# Cytotoxic Profiling of Annotated and Diverse Chemical Libraries Using Quantitative High-Throughput Screening

**DOI:** 10.1101/404665

**Authors:** Olivia W. Lee, Shelley Austin, Madison Gamma, Dorian M. Cheff, Tobie D. Lee, Kelli M. Wilson, Joseph Johnson, Jameson Travers, John C. Braisted, Rajarshi Guha, Carleen Klumpp-Thomas, Min Shen, Matthew D. Hall

## Abstract

Cell-based phenotypic screening is a commonly used approach to discover biological pathways, novel drug targets, chemical probes and high-quality hit-to-lead molecules. Many hits identified from high-throughput screening campaigns are ruled out through a series of follow-up potency, selectivity/specificity, and cytotoxicity assays. Prioritization of molecules with little or no cytotoxicity for downstream evaluation can influence the future direction of projects, so cytotoxicity profiling of screening libraries at an early stage is essential for increasing the likelihood of candidate success. In this study, we assessed cell-based cytotoxicity of nearly 10,000 compounds in NCATS annotated libraries, and over 100,000 compounds in a diversity library, against four ‘normal’ cell lines (HEK 293, NIH 3T3, CRL-7250 and HaCat) and one cancer cell line (KB 3-1, a HeLa subline). This large-scale library profiling was analyzed for overall screening outcomes, hit rates, pan-activity and selectivity. For the annotated library, we also examined the primary targets and mechanistic pathways regularly associated with cell death. To our knowledge, this is the first study to use high-throughput screening to profile a large screening collection (>100,000 compounds) for cytotoxicity in both normal and cancer cell lines. The results generated here constitutes a valuable resource for the scientific community and provides insight on the extent of cytotoxic compounds in screening libraries, identifying and avoiding compounds with cytotoxicity during high-throughput screening campaigns.

## Introduction

The development of new chemical probes enables therapeutic target validation, hypothesis testing, and new insight into the biological role of genes and proteins (1). At the NIH National Center for Advancing Translational Sciences (NCATS), the development of small molecule probes for the scientific community allows understanding of rare and neglected diseases, novel targets, and enables basic biological understanding of the “undrugged” genome. This is accomplished through a team science approach that begins with assay development and automated quantitative high-throughput screening (qHTS) with a small molecule library to identify active hits (2). One or more chemotypes that emerge in a qHTS campaign may progress to medicinal chemistry to develop a small molecule probe with strong biochemical and/or cell-based activity, specificity, and optimized properties to enable use in *in vivo* models. To accommodate unbiased qHTS discovery for medicinal chemistry, large libraries of small molecules are created and curated, containing molecules that capture diverse chemical space that are ideally synthetically tractable. These large libraries of small molecules are generally referred to ‘diversity’ libraries or collections (3).

A second significant discovery screening strategy involves the creation of libraries of ‘annotated’ small molecules. Annotated libraries contain drugs, probes and tool molecules with one or more known mechanisms of action (4). They have emerged as information-rich databases to integrate both biological and chemical data. These can be screened in (primarily) cell or organism-based assays to identify targets relevant to a phenotype, or for potential drug repurposing (5). While compounds in a diversity library are expected to demonstrate weak biological activity, annotated libraries by definition are medicinal chemistry-optimized products with known activity and in many cases, known mechanism of actions. Both diversity collection and annotated library screening are important components of the NCATS Chemical Genomics Center (NCGC) program (6).

Throughout a qHTS campaign, the activities of hits are confirmed in a re-test, and a number of orthogonal and counter-assays are performed to confirm that the observed modulatory activity of is ‘on-target’. This is also to ensure that compounds demonstrating artefactual activity in an assay are triaged. For example, biochemical and cell-based assays that utilize firefly luciferase (fLuc) are sensitive to compounds that modulate luciferase activity (7). To ensure this is not the case, the screening libraries at NCATS are ‘profiled’ for inhibitory activity of fLuc, allowing compounds that interfere with luciferase to be automatically triaged from hit lists without the need for additional screening. In one study it was shown that ∼5% of compounds in a qHTS library inhibit fLuc (8). Perhaps the most notorious example of the need to perform counter-assays is the drug ataluren (PTC124) approved for treatment of patients with Duchenne muscular dystrophy, that may have been discovered due to fLuc inhibition rather than on-target activity (9).

A resurgence in cell-based screening, both target-based and phenotypic, means that an increasing number of cell-based qHTS campaigns are performed. Yet the profiling of libraries against cell-based assay read-outs is rarely reported. An early report of assessing compound cytotoxicity across a library of 1,408 compounds appears in the literature (10), and more recently, two studies examining cell killing with multiplexed assays were reported; one using a high-content assay against ∼12,000 screening library (‘diversity’) compounds (11), and the other using a multiplex assay against ∼10,000 environmental toxins (12). A significant collection of disease-agnostic ‘annotated’ small molecules exist in libraries at NCATS, including a collection of drugs approved by American, European and Japanese therapeutic regulatory agencies (the ‘NCATS Pharmaceutical Collection’, NPC (13)), and many small molecules reported as tools, probes, or clinical/pre-clinical candidates. Understandably, a significant proportion of these molecules arose from oncology programs, and a portion of NCATS’ collaborations are oncology-related phenotypic screens. We were motivated to profile the activity of our annotated libraries for two reasons: first, to allow scientists to have a reference dataset for discerning compounds whose activity is ‘selective’ for cancer cell lines versus a set of non-cancer ‘normal’ cell lines; and second, to enable scientists to discriminate promiscuous/cytotoxic compounds when reviewing data from cell-based/phenotypic assays and provide valuable input for prioritizing compounds for further evaluation.

To this end, we assessed the cytotoxicity of NCATS annotated libraries of nearly 10,000 compounds against four ‘normal’ cell lines (HEK 293, NIH 3T3, CRL-7250 and HaCat) and a cancer cell line (KB 3-1, a HeLa subline), and examined the hit rates and mechanistic pathways regularly associated with cell death. Furthermore, we assessed that activity of a diversity library (>100,000 compounds) against two ‘normal’ cell lines (HEK 293 and NIH 3T3), assessed hit rate, and compared active compounds against a cancer cell line (KB 3-1). This study provides insight into the extent of cell-based killing activity in annotated and diversity libraries, and the importance of confirming that active compounds in phenotypic screens are not cytotoxic.

## Materials and Methods

### Profiling annotated/diversity libraries and cherry-picked compounds

HEK 293, NIH 3T3, CRL-7250, HACAT and KB 3-1 cells were seeded into white 1536-well plates using a Multidrop Combi peristaltic dispenser (ThermoFisher, Waltham, MA) at a density of 250, 400, 500, 500, 500 cells/well in 5 μL of medium respectively. A pintool (Kalypsys) was used to transfer 23 nL of compound solution to the 1536-well assay plates. After 48 hr incubation at 37 °C, 5% CO and 85% humidity, 2.5 μL of CellTiter-Glo (Promega) was dispensed into each well using a dispenser (Aspect Automation, St. Paul, MN) with solenoid valves (Lee Valves, Westbrook CT). Plates were left at room temperature for 10 min before imaging the ATP-coupled luminescence using a ViewLux microplate imager (PerkinElmer, Waltham, MA).

### Luciferase assay protocol

Assays determining firefly luciferase inhibition were performed as previously described (14). Briefly, 3 μL of luciferase substrate solution (10 μM ATP, 10 μM D-Luciferin, 10 mM Mg-Acetate, 0.01% Tween-20, 0.05% BSA, 50 mM Tris Acetate, pH 7.6, in final 4 μL volume) was dispensed into each well of white, solid bottom, 1536-well plates using a dispenser. A pintool (Kalypsis) was used to transfer 23 nL of compound solution to the assay plates. Following a 15 minute incubation at room temperature protected from light, 1 μL of purified luciferase enzyme solution was added to a final concentration of 10 nM *Photinus pyralis* luciferase (Sigma). Luminescence was detected by a Viewlux (Perkin-Elmer, Waltham, MA) by using a 10 s exposure time and 2X binning.

### Data analysis and clustering of compounds by activity outcomes

To determine compound activity in the qHTS assay, the concentration-response data for each sample was plotted and modeled by a four parameter logistic fit yielding EC_50_ and efficacy (maximal response) values as previously described (15). Raw plate reads for each titration point were first normalized relative to positive control (2 mM Bortezomib, −100% activity, full inhibition) and DMSO only wells (basal, 0% activity). Data normalization and curve fitting were performed using in-house informatics tools. Compounds were designated as Class 1–4 according to the type of concentration–response curve (CRC) observed. In brief, Class −1.1 and −1.2 were the highest-confidence complete CRCs containing upper and lower asymptotes with efficacies ≥ 80% and < 80%, respectively. Class −2.1 and −2.2 were incomplete CRCs having only one asymptote with efficacy ≥ 80% and < 80%, respectively. Class −3 CRCs showed activity at only the highest concentration or were poorly fit. Class 4 CRCs were inactive having a curve-fit of insufficient efficacy or lacking a fit altogether.

Compounds were further clustered hierarchically using TIBCO Spotfire 6.0.0 (Spotfire Inc., Cambridge, MA. https://spotfire.tibco.com/) based on their activity outcomes from the primary or follow up screen across different cell lines. Compound’s AUC calculated based on the qHTS data analysis and curve fittings were utilized for clustering. In the heatmap, darker color indicates compounds that are more potent and efficacious, i.e. high-quality actives, and lighter color indicates less potent and efficacious compounds. If a compound didn’t show any activity in an assay, it was highlighted as white in the heatmap.

### Statistical analysis

In order to determine whether the hits predominantly identified from the specific therapeutic categories were over-represented in the chemical library, an enrichment analysis was implemented against the drug library. 9,893 compounds in the annotated library were broken down to different therapeutic categories based on their primary mechanisms of action and pharmaceutical indications, the enrichment was calculated from the following formula: E = a/n, given a is the number of actives and n is the total number of drugs in each therapeutic category. Fisher’s exact tested was used as a measure of the consensus cytotoxicity potential of compounds in each MOA; all calculations were performed in R statistical computing software (https://www.r-project.org/).

## Results

### Profiling of an annotated library

Annotated libraries were profiled for cell viability by performing a primary screen against four normal cell lines using CellTiter-Glo (CTG) as the assay read-out (screening assay protocol displayed in **Table 1**). Three of the cell lines are immortalized (HEK 293, NIH 3T3 and HaCat), while CRL-7250 is a primary cell line. A total of 9,893 compounds were tested in either an 8-pt concentration-response ranging from 0.6 nM to 46 µM (1:5 dilution) or an 11-pt concentration-response ranging from 0.8 nM to 46 µM (1:3 dilution) according to different compound plating mechanisms.

**Table 1.**
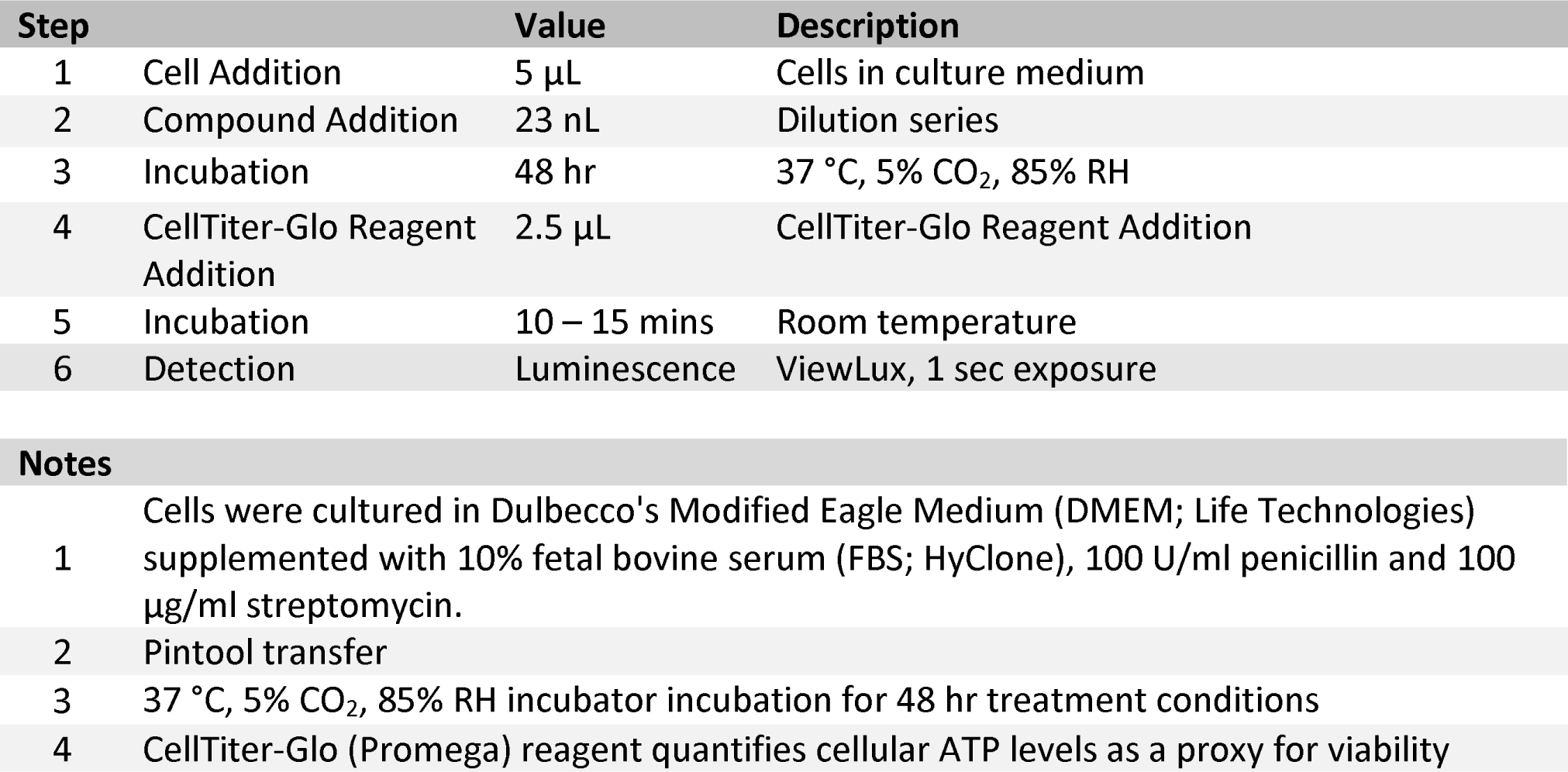
CellTiter-Glo assay protocol

| Step |  | Value | Description |
| --- | --- | --- | --- |
| 1 | Cell Addition | 5 μL | Cells in culture medium |
| 2 | Compound Addition | 23 nL | Dilution series |
| 3 | Incubation | 48 hr | 37 °C, 5% CO<sub>2</sub>, 85% RH |
| 4 | CellTiter-Glo Reagent Addition | 2.5 μL | CellTiter-Glo Reagent Addition |
| 5 | Incubation | 10 – 15 mins | Room temperature |
| 6 | Detection | Luminescence | ViewLux, 1 sec exposure |

**Notes**
- Cells were cultured in Dulbecco's Modified Eagle Medium (DMEM; Life Technologies) supplemented with 10% fetal bovine serum (FBS; HyClone), 100 U/ml penicillin and 100 $\mu$ g/ml streptomycin. - Pintool transfer - 37 °C, 5% CO<sub>2</sub>, 85% RH incubator incubation for 48 hr treatment conditions - CellTiter-Glo (Promega) reagent quantifies cellular ATP levels as a proxy for viability

Variation of sensitivity was observed across all four cell lines as shown in **Table 2**. The HEK 293 cell line was more sensitive to compound treatment (**Figure 1a**), with 41% of compounds eliciting a dose-response cell killing effect (16% high-quality actives, 25% low-quality actives), while the three remaining cell lines demonstrated reduced sensitivity (7.9–10.6% high-quality actives, 14.2-18.1% low-quality actives), where the ‘high-quality’ actives were defined as compounds in class −1.1, −1.2, −2.1, and −2.2, with maximal response higher than 50% and EC_50_ values less than 10 µM. The detailed information for data analysis and curve class definition is provided in Methods section. Comparison of compounds active against the four cell lines (**Figure 1b**, Venn diagram) revealed 496 compounds active only against HEK 293 cells (consistent with its greater sensitivity to compounds). 409 compounds demonstrated pan-activity against all four cell lines, which corresponds to a consensus hit rate of 4.1% for the entire annotated library.

**Figure 1.**
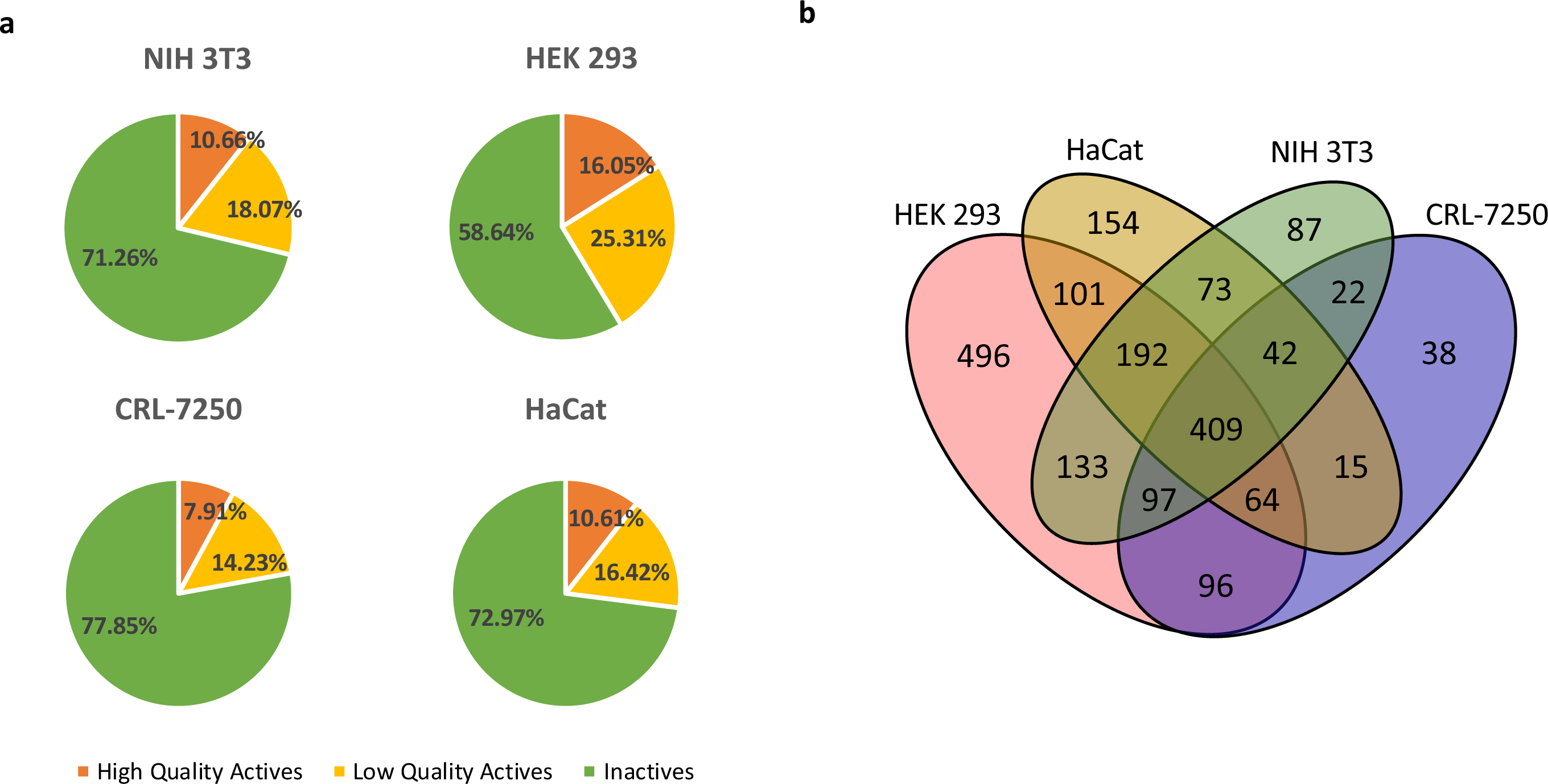
**a)** Pie chart distribution of high-quality actives (orange), low-quality actives (yellow), and inactives (green) identified from the annotated compound library qHTS against the four normal cell lines for 48 hr incubation condition. High-quality actives: compounds in class −1.1, −1.2, −2.1, and −2.2, maximum response ≤ −50%, and EC_50_ value ≤ 10 μM. Inactives: compounds in curve class 4; low-quality actives: compounds with shallow curves, single-point activity or inconclusive activity. **b)** Compound overlapping Venn diagram for HEK 293, HaCat, NIH 3T3, and CRL-7250 cell lines. Number of high-quality actives in each cell line and number of compounds overlapped were calculated.

**Table 2.** Summary table of qHTS screen against the annotated library in normal cell lines.

| Cell line | Activity category | Number of compounds |
| --- | --- | --- |
| HEK 293 | High-quality actives* | 1588 |
|  | Low-quality actives* | 2504 |
|  | Inactives* | 5801 |
| NIH 3T3 | High-quality actives | 1055 |
|  | Low-quality actives | 1788 |
|  | Inactives | 7050 |
| CRL-7250 | High-quality actives | 783 |
|  | Low-quality actives | 1408 |
|  | Inactives | 7702 |
| HaCat | High-quality actives | 1050 |
|  | Low-quality actives | 1624 |
|  | Inactives | 7219 |
| Total number of compounds in Annotated library |  | 9893 |

\*High-quality active: compounds in class -1.1, -1.2, -2.1, and -2.2, maximum response ≤ -50%, and EC<sub>50</sub> value ≤ 10 μM; inactives: compounds in curve class 4; low-quality actives: compounds with shallow curves, single-point activity or inconclusive activity.

Comparison of the AUC (Area Under the dose-response Curve) values of each compound using unbiased hierarchical clustering across four cell lines revealed some cross-correlation of activity, with the HEK 293 cells clustering away from the other cell lines (**Figure 2a**). Given the long-term intention of utilizing profiling data to make comparisons with activity against cancer cell lines, we also tested the library against KB 3-1 human adenocarcinoma cells (KB 3-1 cells are a HeLa sub-clone). Intriguingly, the KB 3-1 cell line did not demonstrate remarkably different sensitivity compared to the normal cells, but surprisingly its sensitivity clustered close with the NIH 3T3 murine fibroblast line. The pair-wise AUC correlation with R^2^ for all actives among five cell lines were calculated and shown in **Figure 2b**, the R^2^ ranges from 0.31 to 0.77. NIH 3T3 and KB 3-1 showed the strongest correlation with R^2^ = 0.77. Violin plots of the distribution of sensitivity to compounds across the libraries reinforce that the majority of compounds were not cytotoxic in cell lines (**Figure 2c**, pink area), while focused plots of only high-quality actives (**Figure 2c**, cyan area) show that only a small number of compounds are highly cytotoxic (AUC < −300) against cell lines.

**Figure 2.**
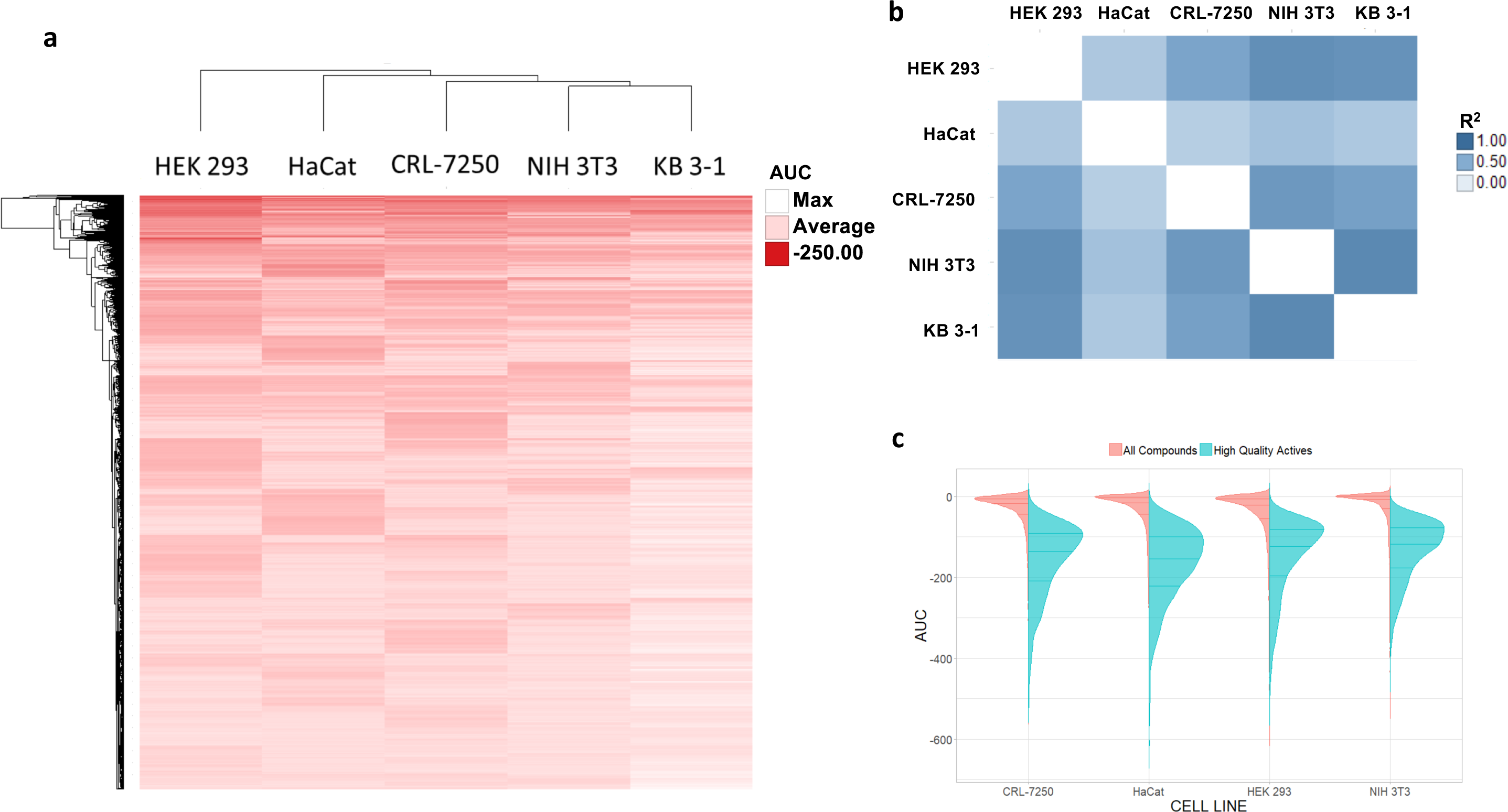
**a)** The comparison of AUC values of each compound in four normal cell lines and KB 3-1 human adenocarcinoma cells. In the heat map each row corresponds to a compound and each column to a cell line. Darker red color indicates more potent and efficacious compound. **b)** Pair-wise R^2^ correlation matrix among five cell lines. Only high-quality actives were included in this analysis and the Pearson correlation was based on AUC values. **c)** Split violin plot showing the distribution of AUC values for all compounds screened and high-quality actives across four normal cell lines. The lines within each distribution area represent the 0.25, 0.5 and 0.75 quantiles.

One strength of annotated compound libraries is the ability to integrate compound mechanism of action (MOA) and targets into downstream analysis. Each compound that met the criteria for high-quality active in each cell line were aggregated by mechanism of action. The treemap shown in **Figure 3** contains 1 box for each MOA containing three or more compounds. The size of each box is representative of the number of compounds with that MOA. The MOAs with the most compounds were antibacterial agents and PI3 Kinase inhibitors with 42 compounds present in each. We also assessed average AUC for each MOA which is represented in **Figure 3** by the color gradient for each box. Kinesin like Spindle Protein (KIF11) inhibitors and proteasome inhibitors had the lowest average AUC of all MOAs.

**Figure 3.**
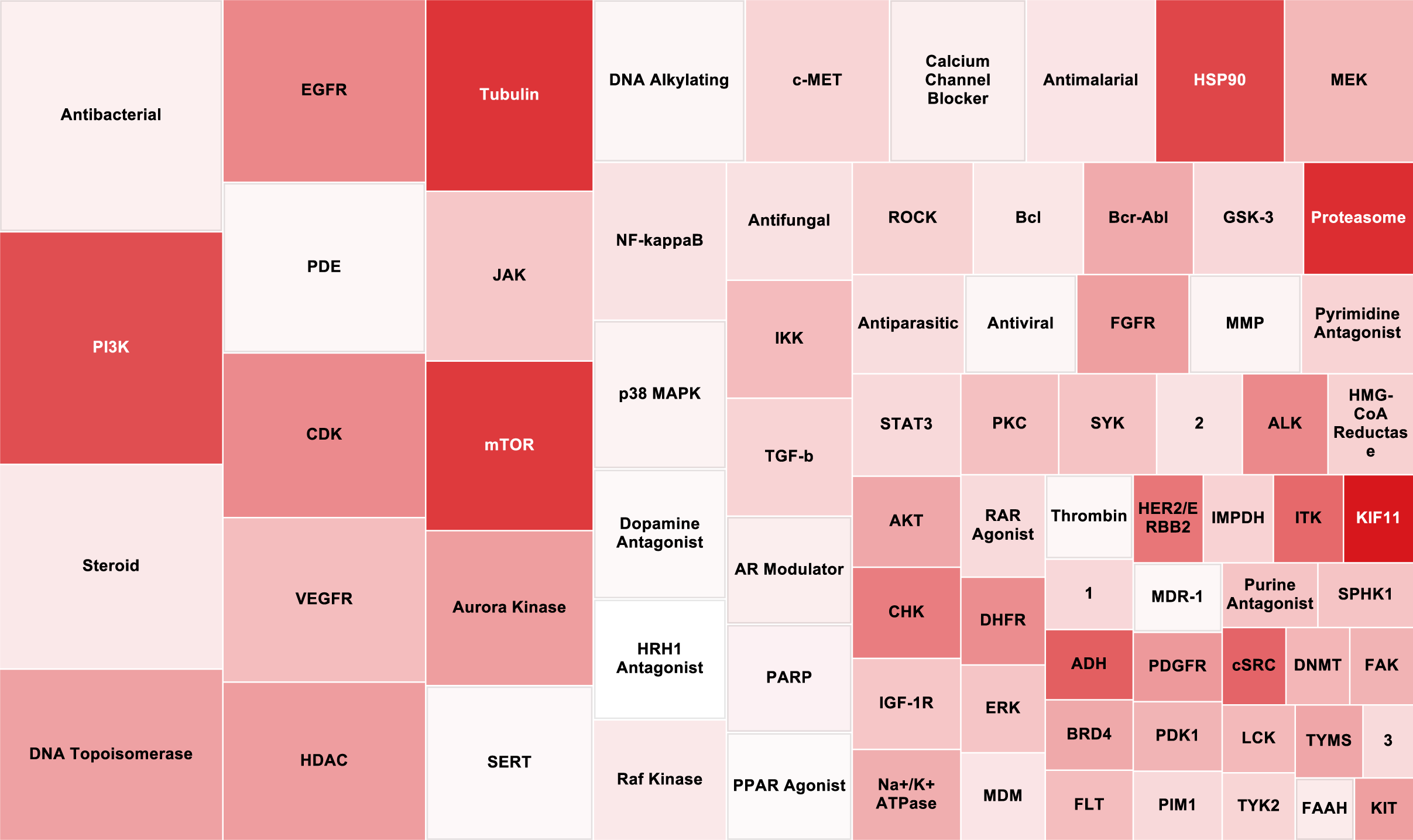
A treemap representation of the mechanisms of action (MOA) of all high-quality actives from the annotated library compounds screened. Box size represents the total number of compounds representing each MOA (bigger box size indicates more compounds present in the high-quality actives). Color represents the average AUC from the cytotoxicity screen in four normal cell lines (darker red indicates a lower AUC meaning a more potent and efficacious hits). 1 = Lineage specific differentiation; 2 = RNA polymerase; 3 = Immuno-suppressant.

There were several instances where multiple molecules with the same target were screened. To examine which MOAs were enriched among molecules active against cell lines, the proportion of active drugs to total drugs for each MOA target were used to derive an ‘enrichment ratio’ (**Figure 4a**, where ratio = 1 represents 100% active, ratio = 0 represents none active). All classes of MOA targets with at least five compounds presented in the annotated library were included in the enrichment analysis. The top five target classes that resulted in consensus cell killing against the normal cell lines were proteasome, heat shock protein 90 (HSP90), anaplastic lymphoma kinase (ALK), mammalian target of rapamycin (mTOR), and cyclin-dependent kinase (CDK), with the enrichment ratio greater than 50%. To examine the significance of the association between consensus activity outcomes with the primary MOA, Fisher’s exact test was applied on the entire data set and the statistical significance was reported in **Supplementary Table 1**. Eight out of eleven highly enriched MOA classes have P-value < 0.05, and seven classes with P-value < 0.001, indicating that compounds with similar activity profiles as determined in normal cell line screens tend to share similar annotated MOAs. These highly enriched MOAs (along with many other targets shown in **Figure 4a**) are generally associated with anti-cancer cell killing by targeting essential cellular processes. For example, nine of ten proteasome inhibitors tested resulted in acute cytotoxicity across all four cell lines (see proteasome inhibitor delanzomib, **Figure 4b**). As a reference point, the cancer cell line KB 3-1 did not demonstrate hypersensitivity compared with the normal cell lines, this is also consistent to the results we discussed in the heatmap clustering analysis (**Figure 2a**). The global cytotoxicity of proteasome inhibitors is displayed by radar plot (**Figure 4c**) and is extended to other mechanisms of actions (**supplementary Figure 1**).

**Figure 4.**
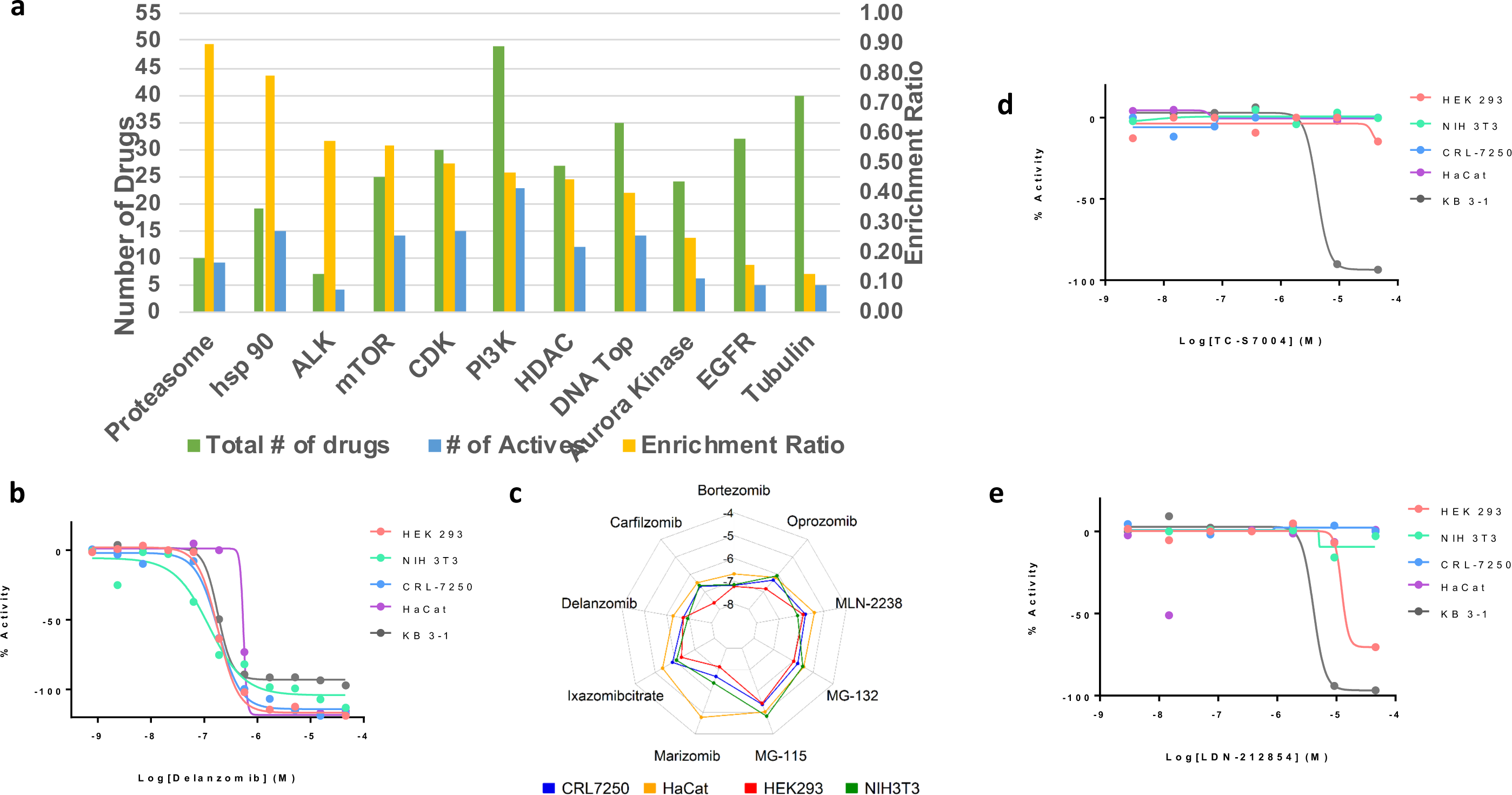
**a)** Enrichment analysis of active agents in each drug category. Enrichment ratio = the number of actives/the total number of drugs in each drug category. **b)** Dose-response curves for delanzomib, a representative proteasome inhibitor, in cytotoxicity screens. **c)** A radar plot displaying Log EC_50_ of all active proteasome inhibitors in four normal cell lines. **d)** Dose-response curves for TC-S7004, a potent and selective DYRK1A/B inhibitor, in cytotoxicity screens, including KB 3-1 cells. **e)** Dose-response curves for LDN-212854, an ALK2 inhibitor, in cytotoxicity screens, including KB 3-1 cells.

As outlined, one use of cell profiling data for mechanistically-annotated screening libraries is to enable the identification of targeted agents selectively killing cancer cells. Here we defined that a selective agent should show strong cell killing to KB 3-1 cells with EC_50_ ≤ 10 µM and have no effect to normal cell lines or at least 10-fold of EC_50_ shift in normal cell lines. Two such examples of selective killing are displayed in **Figures 4d** and **4e**. First, the dual specificity tyrosine phosphorylation regulated kinase 1A and 1B (DYRK1A and DYR1B) inhibitor TC-S7004 demonstrated complete killing of KB 3-1 cells (EC_50_ = 3.2 µM), with no effect on any of the four normal cell lines. Second, the potent and selective BMP receptor inhibitor LDN-212854 demonstrated complete killing of KB 3-1 cells (EC_50_ = 4.1 µM), with no effect on NIH 3T3 and CRL-7250 cells, and only partial killing of one normal line (HEK 293, EC_50_ = 16.4 µM).

Data for annotated library screen was deposited in PubChem with AID 1296008, and you can access this assay with the following link: https://pubchem.ncbi.nlm.nih.gov/assay/assay.cgi?aid=1296008

### Profiling of a diversity library

The diversity library was profiled for cell viability by performing a primary screen against HEK 293 and NIH 3T3 cell lines, using CTG as the assay read-out (screening assay protocol displayed in **Table 1**). A total of 102,726 compounds were tested in a 4-pt dose response manner at varied concentrations with nearly 80% of the compounds tested at 115, 58, 12 and 2.3 µM. 30.0% and 34.6% of the library was cytotoxic to cell lines NIH 3T3 and HEK 293 respectively, with 1.2% of the library classified as ‘high-quality’ actives in each of the cell lines (**Figures 5a and 5b**). Of these high-quality actives, there were 285 compounds overlapped and shared similar activity in both NIH 3T3 and HEK 293 (**Figure 5 c**).

**Figure 5.**
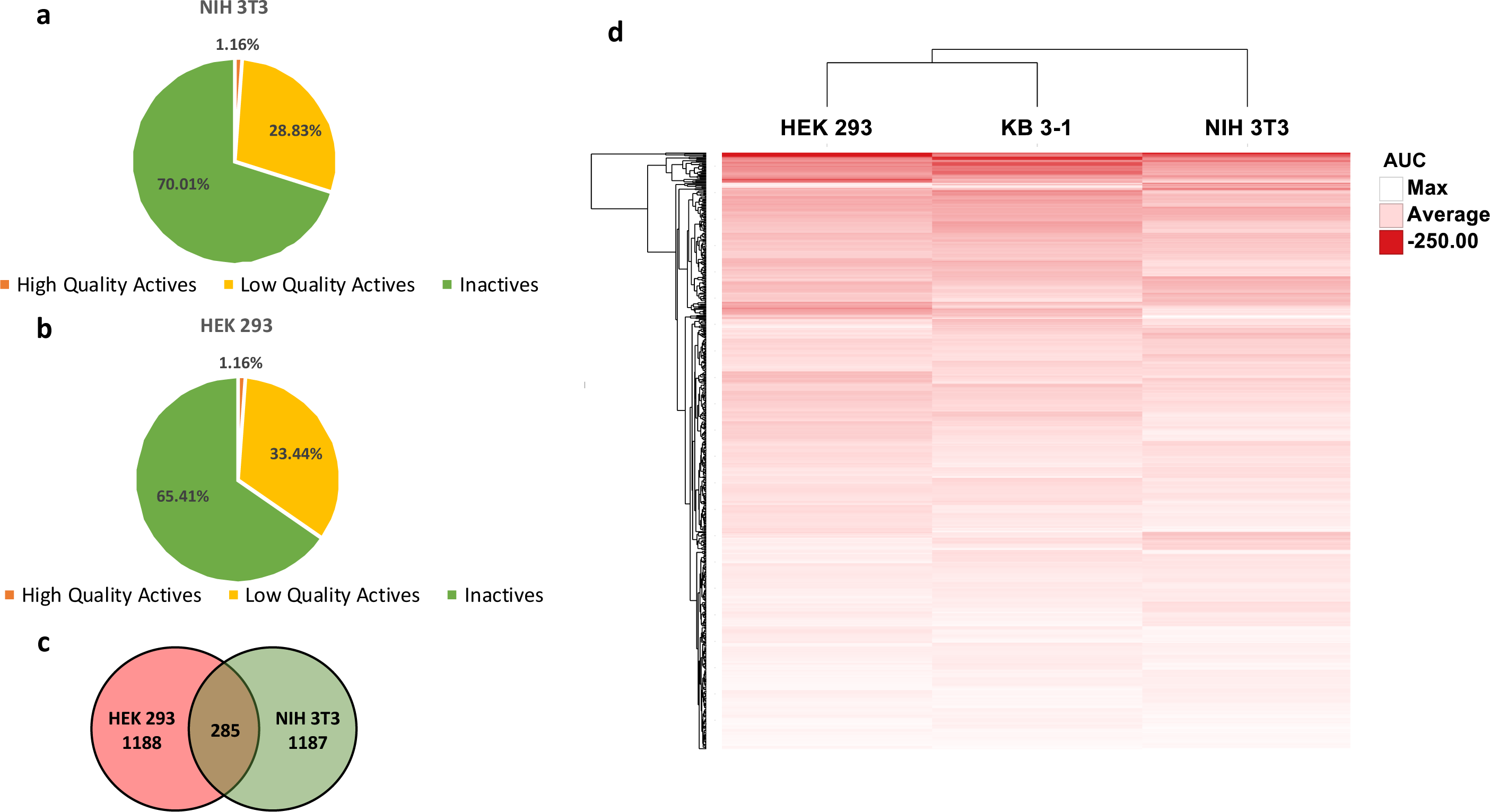
**a-b)** Pie chart distribution of high-quality actives (orange), low-quality actives (yellow), and inactives (green) identified from the qHTS against diversity collection compound library in NIH 3T3 and HEK 293 cell line, respectively. **c)** Compound overlapping Venn diagram for NIH 3T3 and HEK 293 cell lines. Number of high-quality actives in each cell line was calculated and number of compounds overlapped were labeled. **d)** The comparison of AUC values of 588 cherry-picked compounds in HEK 293, NIH 3T3 and KB 3-1 cell lines. In the heat map each row corresponds to a compound and a column to a cell line. Darker red color indicates more potent and efficacious compound.

After cherry-picking 588 hits based on their potency, maximal response in primary qHTS assay and structural features, confirmation of activity was performed in 12 dose-point testing for all the cherry-picked compounds against HEK 293, NIH 3T3, along with an orthogonal test against the KB 3-1 cell line. The final concentration of the compounds in the 5 µl assay volume ranged from 0.3 nM to 46 µM. To analyze the data, we generated a heat map with hierarchical clustering analysis (dendrogram) of compound activities based on their activity outcomes, which shows that NIH 3T3 clusters away from HEK 293 and KB 3-1 (**Figure 5d**).

The Promega CTG reagent utilizes an engineered version of firefly luciferase to measure ATP concentrations. As such, we tested compounds that appeared cytotoxic against recombinant fLuc in a biochemical assay to triage any compounds that were inhibiting fLuc rather than reducing cell viability. A number of compounds inhibited fLuc, and two examples from a novel fLuc inhibitor chemotype are shown in **Figures 6a and 6b**. In both cases, the compounds appeared equally active against all three cell lines in the CTG assay and demonstrated potent fLuc inhibition. The Promega CTG reagent contains a proprietary engineered fLuc that claims to be resistant to inhibition compared with wild-type fLuc (16). This indeed appears to be the case, as a counter-assay testing hits in cell-free media with CTG and ATP added as substrate demonstrated at least a 10-fold lower sensitivity to inhibition. Bright-field microscopy of cells grown in the presence of NCGC00413522 and NCGC00413607 (**Figure 6c**) demonstrated unaffected cell growth at 48 hr at concentrations suggested to be cytotoxic by CTG dose-response data (DMSO and 500 nM bortezomib as negative and positive control respectively).

**Figure 6.**
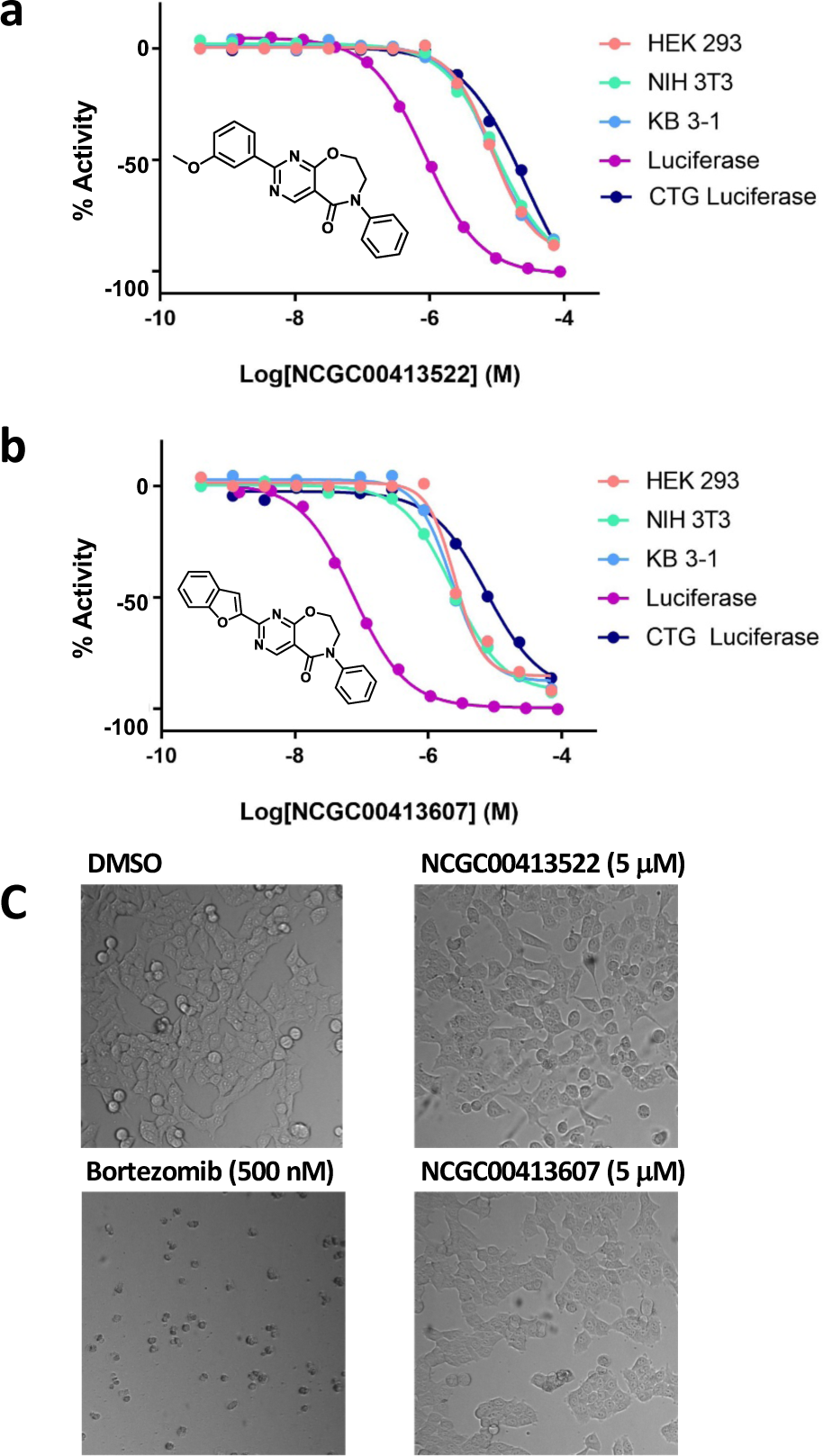
**a-b)** Dose-response curves for NCGC00413522 and NCGC00413607 in cytotoxicity screens, luciferase inhibition assay and CellTiter-Glo luciferase assay. **c)** Representative bright field images of HEK293 cells after 48 hr treatment.

Moreover, comparing the AUC values of high-quality actives in the three cell lines, we found that they moderately correlated: NIH 3T3 vs HEK 293, R^2^ = 0.77; NIH 3T3 vs KB 3-1, R^2^ = 0.59; KB 3-1 vs HEK 293, R^2^ = 0.61 (**Figures 7 a, b, c**). Sixty-one of the top-ranking hits were active across all cell lines (pan-active), whereas others were more selective toward a specific cell line or cell lines. If only taking the pan-actives that showed strong cytotoxicity against all three cell lines in the analysis, it was clearly shown that most of the compounds have very good EC_50_ agreement with low standard deviation, except for few outlier compounds that showed differential cell killing effect in three cell lines (**Figure 7d**). These outlier compounds had greater cell killing against two normal cell lines (HEK 293 and NIH 3T3) although the selectivity window to the cancer cell line KB-3-1 was not significant, individual dose-response curves were shown in **Supplementary Figure 2**. Among these five compounds, NCGC00420512, NCGC00420455 and NCGC00420435 belonged to the same structural chemotype with a tetrahydropyrazolo-pyrimidine core. To examine if the observed selectivity is a chemotype-specific outcome, we further analyzed the screening data through substructure search and found 172 compounds sharing the same tetrahydropyrazolo-pyrimidine scaffold presented in our diverse library, but none of the remaining 169 compounds showed similar differential cell killing effect, indicating that the selectivity is irrelevant to the specific structural features.

**Figure 7.**
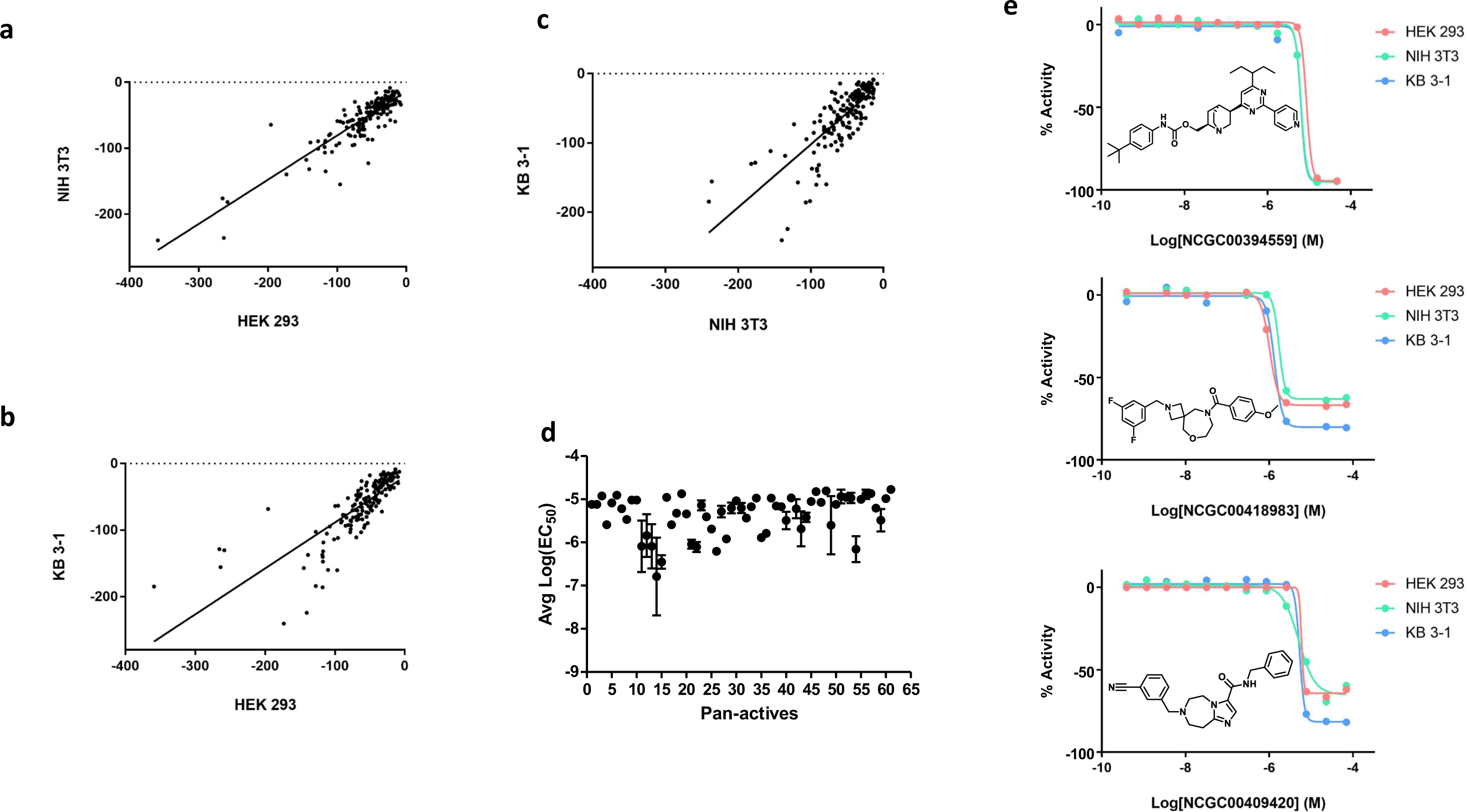
**a-c)** Activity correlation plot between HEK 293, NIH 3T3 and KB 3-1 cell lines. Only high-quality actives were included in this analysis and the correlation was based on AUC values. **d)** Log EC_50_ distribution of the pan-active compounds in all three cell line screens. Error bars represent the standard deviation of Log EC_50_. **e)** Dose-response curves for representative compounds in diversity library showing pan-activity across three cell lines.

The remaining 56 pan-actives were cytotoxic across all three cell lines lacking any selectivity toward either normal or cancer lines. Representatives of the top ranking pan-actives were shown in **Figure 7e**. Pan-actives were further clustered using fingerprint structural keys and Tanimoto coefficient as similarity metric in Molecular Operating Environment software (MOE, https://www.chemcomp.com/). We identified three enriched clusters based on their structural similarity using MACCS fingerprint and Tanimoto coefficient (**Table 3**). The most significant enriched scaffolds are in Clusters 1 and 2. Members in Cluster 1 shared the same pyrimidine-quinuclidine core, with comparable EC_50_ values across all three cell lines ranging from 7 to 12 µM, except one compound NCGC00421344 which was > 10-fold more potent in HEK 293 cell than the other two cell lines. Cluster 2 had the spiro structural feature and it was the most cytotoxic cluster among all four clusters discussed in **Table 3**. Compounds in this cluster showed almost identical activity across three cell lines, with EC_50_ values ranging from 0.5 to 3 µM for most of the compounds. For example, NCGC00419015 was the most cytotoxic agent in Cluster 2 with EC_50_ values for HEK 293, NIH 3T3 and KB 3-1 being 0.51, 0.72 and 0.64 µM, respectively. Furthermore, Cluster 3 with tetrahydro-imidazo-diazepine core was a less enriched cluster but also showed as non-selective cluster in which compounds were lacking selectivity against a specific cell line. We also ran the pan-actives against multiple toxicity datasets including Bursi Mutagenicity dataset (17) (4,337 compounds with mutagenicity data) and National Toxicology Program dataset (18) (503 structures with carcinogenicity data for male/female mouse and male/female rat), but found no overlapping structures which indicated that the chemotypes identified from our diverse library profiling are new additions to the well-known toxic collections.

**Table 3.**
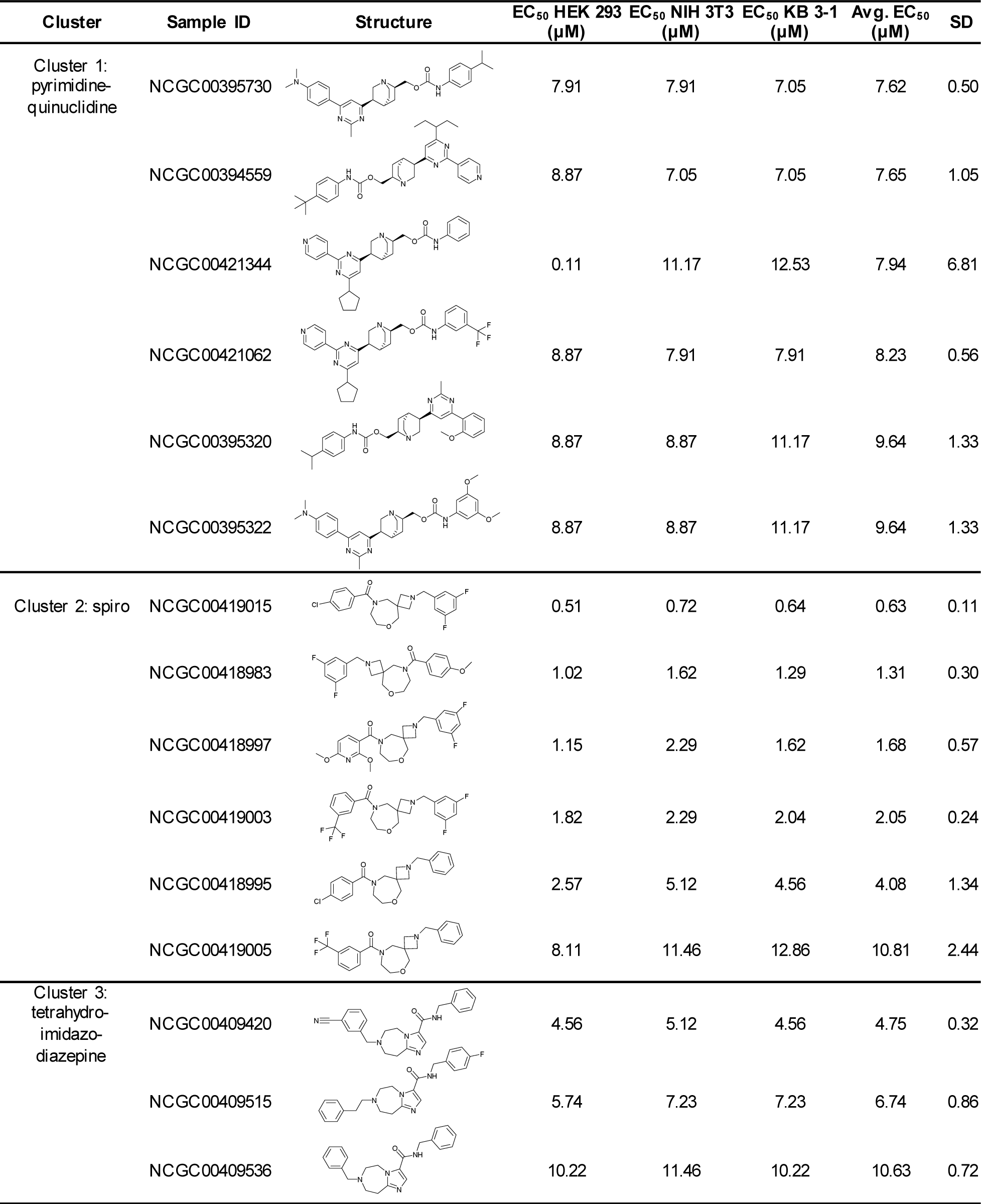
Representative clustered hits identified from qHTS against diversity chemical library.

| Cluster | Sample ID | Structure | EC<sub>50</sub> HEK 293<br>(μM) | EC<sub>50</sub> NIH 3T3<br>(μM) | EC<sub>50</sub> KB 3-1<br>(μM) | Avg. EC<sub>50</sub><br>(μM) | SD |
| --- | --- | --- | --- | --- | --- | --- | --- |
| Cluster 1:<br>pyrimidine-quinuclidine | NCGC00395730 |  | 7.91 | 7.91 | 7.05 | 7.62 | 0.50 |
|  | NCGC00394559 |  | 8.87 | 7.05 | 7.05 | 7.65 | 1.05 |
|  | NCGC00421344 |  | 0.11 | 11.17 | 12.53 | 7.94 | 6.81 |
|  | NCGC00421062 |  | 8.87 | 7.91 | 7.91 | 8.23 | 0.56 |
|  | NCGC00395320 |  | 8.87 | 8.87 | 11.17 | 9.64 | 1.33 |
|  | NCGC00395322 |  | 8.87 | 8.87 | 11.17 | 9.64 | 1.33 |
| Cluster 2: spiro | NCGC00419015 |  | 0.51 | 0.72 | 0.64 | 0.63 | 0.11 |
|  | NCGC00418983 |  | 1.02 | 1.62 | 1.29 | 1.31 | 0.30 |
|  | NCGC00418997 |  | 1.15 | 2.29 | 1.62 | 1.68 | 0.57 |
|  | NCGC00419003 |  | 1.82 | 2.29 | 2.04 | 2.05 | 0.24 |
|  | NCGC00418995 |  | 2.57 | 5.12 | 4.56 | 4.08 | 1.34 |
|  | NCGC00419005 |  | 8.11 | 11.46 | 12.86 | 10.81 | 2.44 |
| Cluster 3:<br>tetrahydro-imidazo-diazepine | NCGC00409420 |  | 4.56 | 5.12 | 4.56 | 4.75 | 0.32 |
|  | NCGC00409515 |  | 5.74 | 7.23 | 7.23 | 6.74 | 0.86 |
|  | NCGC00409536 |  | 10.22 | 11.46 | 10.22 | 10.63 | 0.72 |

Data for diverse chemical library screen was deposited in PubChem with AID 1345083 for HEK 293 cell line and AID 1345082 for NIH 3T3 cell line. You can access the assay data with the following links:

https://pubchem.ncbi.nlm.nih.gov/assay/assay.cgi?aid=1345083 https://pubchem.ncbi.nlm.nih.gov/assay/assay.cgi?aid=1345082

## Discussion

Here, we describe our effort to profile the cytotoxicity of two distinct libraries: 1. our annotated libraries (nearly 10,000 compounds of known MOAs or therapeutic indications) against four ‘normal’ cell lines (HEK 293, NIH 3T3, CRL-7250 and HaCat) and a cancer cell line (KB 3-1, a HeLa sub-line); and 2. a diversity library (>100,000 compounds) against two ‘normal’ cell lines (HEK 293 and NIH 3T3) and a cancer cell line (KB 3-1, a HeLa sub-line). The assays were all performed in quantitative HTS format. Hit rates and mechanistic pathways regularly associated with cell death are described. As one might reasonably anticipate, ‘annotated’ (or mechanistic) libraries containing small molecules developed against targets or known to possess phenotypic activity have a high hit rate (7.91-16.05%) against the cell lines, especially for many annotated molecules were developed against oncology targets.

The large diversity collection revealed a low rate of profound cytotoxicity (1.2%), and this is consistent with the fact that the majority are not optimized against targets, but reinforces the need to confirm toxicity of such compounds through counter-assays as part of screening for compound prioritization and triage. In our case, this profiling data is now used at NCATS to allow for rapid cross-referencing of active compounds in a screen against our dataset of general cytotoxicity in normal cell lines. To facilitate the use of these data by the broader research community, we have also made a significant amount of this data available through PubChem.

What is a ‘normal’ cell line, and what are their limitations? The four ‘normal’ cell lines used in this study were selected for a number of reasons. HEK 293 and NIH 3T3 cell lines are very commonly employed in the scientific literature as ‘control’, ‘normal’, or ‘comparator’ cell lines, along with conventional research uses. A common criticism of these lines is how normal they are given their capacity for uncontrolled cell growth. The HEK 293 cell line is one of the most utilized tool cell lines. It is a neuronal lineage line, generated in 1973 from embryonic kidney cells from an aborted fetus immortalized by transformation with adenovirus (19). The NIH 3T3 murine fibroblast cell line was generated in 1962 from Swiss albino mouse embryo, and spontaneously immortalized after multiple passages (20). The HaCat human keratinocyte cell line spontaneously immortalized in culture and was first reported in 1988 (21). All three of the aforementioned immortalized cell lines are adherent, and all are non-diploid despite being non-cancer non-tumorigenic cells. The CRL-7250 cell line differs from the others in that it is a primary, non-immortalized fibroblast cell line generated from human foreskin (22). The HEK 293 cell line demonstrated greater sensitivity than the other three ‘normal’ cell lines in the annotated library screen.

One potential use for normal cell line profiling data is to enable the identification of compounds selectively active against cancer cell lines based on their target biology. A well-known example are inhibitors of MAPK/ERK kinase (MEK), that elicit acute cell killing in cells harboring somatic activating mutations of Ras, but not in cells expressing wild-type Ras (such as the ‘normal’ cells discussed here (23)). As a proof-of-concept comparison, we tested the KB 3-1 adenocarcinoma cells against the annotated libraries. KB 3-1 cells are a sub-clone of HeLa cells (24), originally called KB squamous cell carcinoma before it was identified as a HeLa contaminant (25)and has subsequently been studied and acknowledged as such. KB cells do not possess activating Ras mutations, but selective activity was seen by a DYRK1A/B inhibitor that killed KB 3-1 cells without affecting the normal cells, demonstrating the utility. This profiling data has already been applied to analysis of multiple oncology-related screens across NCATS.

Perhaps the key result (reassuring from a screening perspective) from cytotoxicity profiling of >100,000 diversity library compounds was the very low rate of cytotoxicity observed, though some chemotype-related activity was observed. Cell lines like HEK 293 (along with cell lines such as Chinese hamster ovary, CHO, cell line) are commonly used to engineer reporter cell lines for cell target-based and high-content assays, and the relative insensitivity to diversity libraries supports their utilization. The only prior study we have identified (also from NCATS but not involving any of the current authors) tested 1,408 compounds against thirteen cell lines, including a number of cancer cell lines along with HEK 293 and NIH 3T3 cells (10). While the compound number was relatively modest in scale, the normal HEK 293 and NIH 3T3 cells were reported to be among the least sensitive, supporting the basis for utilization of normal cell lines rather than cancer cell lines for designing cell-based assays for discovery screening.

The data reported here is made available through PubChem, and it is hoped this data acts as a general guide for normal cell line sensitivity to killing and assists in guiding others in the design of counter-assays for high-throughput screening of cell-based assays.

## Acknowledgements

We thank Dr. Aleksandra Michaloa for advice regarding data analysis. This work was supported by the NCATS Division of Pre-Clinical Innovation Intramural Program.

---

**Supplementary Table 1.** Fisher’s exact test for enriched MOA clusters in compounds showing consensus cell killing.

| **MOA Cluster** | **Number of**<br>**actives in**<br>**each MOA** | **Total**<br>**number**<br>**of actives** | **Total number**<br>**of compounds**<br>**in each MOA** | **Total number of**<br>**compounds with**<br>**MOA** | **P-value** |
| --- | --- | --- | --- | --- | --- |
| Proteasome | 9 | 409 | 10 | 2165 | 5.08E-07 |
| hsp 90 | 15 | 409 | 19 | 2165 | 1.67E-09 |
| ALK | 4 | 409 | 7 | 2165 | 1.47E-02 |
| mTOR | 14 | 409 | 25 | 2165 | 4.26E-06 |
| CDK | 15 | 409 | 30 | 2165 | 1.22E-05 |
| PI3K | 23 | 409 | 49 | 2165 | 2.30E-07 |
| HDAC | 12 | 409 | 27 | 2165 | 3.86E-04 |
| DNA Top | 14 | 409 | 35 | 2165 | 4.78E-04 |
| Aurora Kinase | 6 | 409 | 24 | 2165 | 1.69E-01 |
| EGFR | 5 | 409 | 32 | 2165 | 5.92E-01 |
| Tubulin | 5 | 409 | 40 | 2165 | 7.86E-01 |

**Supplementary Figure 1.**
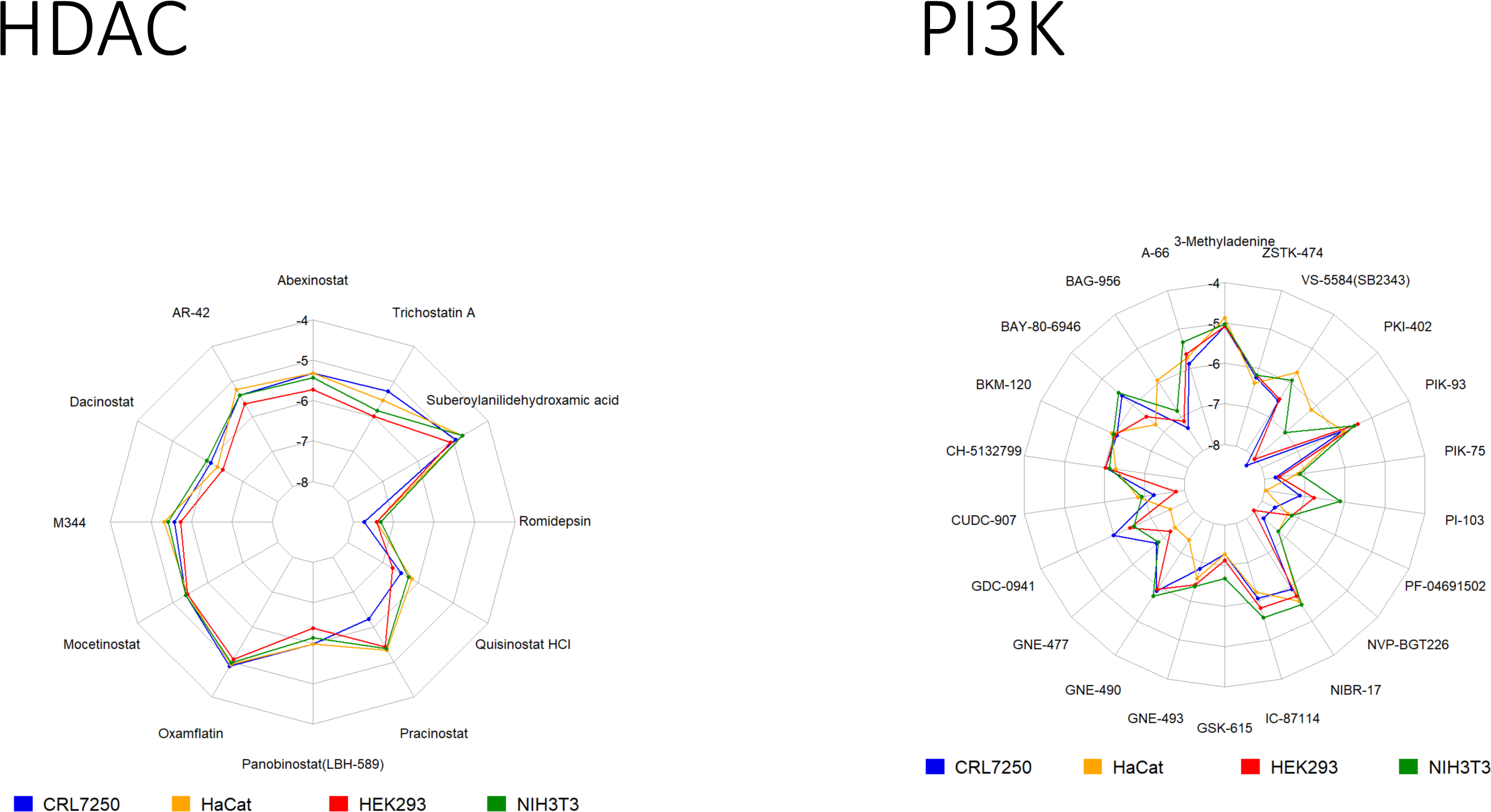
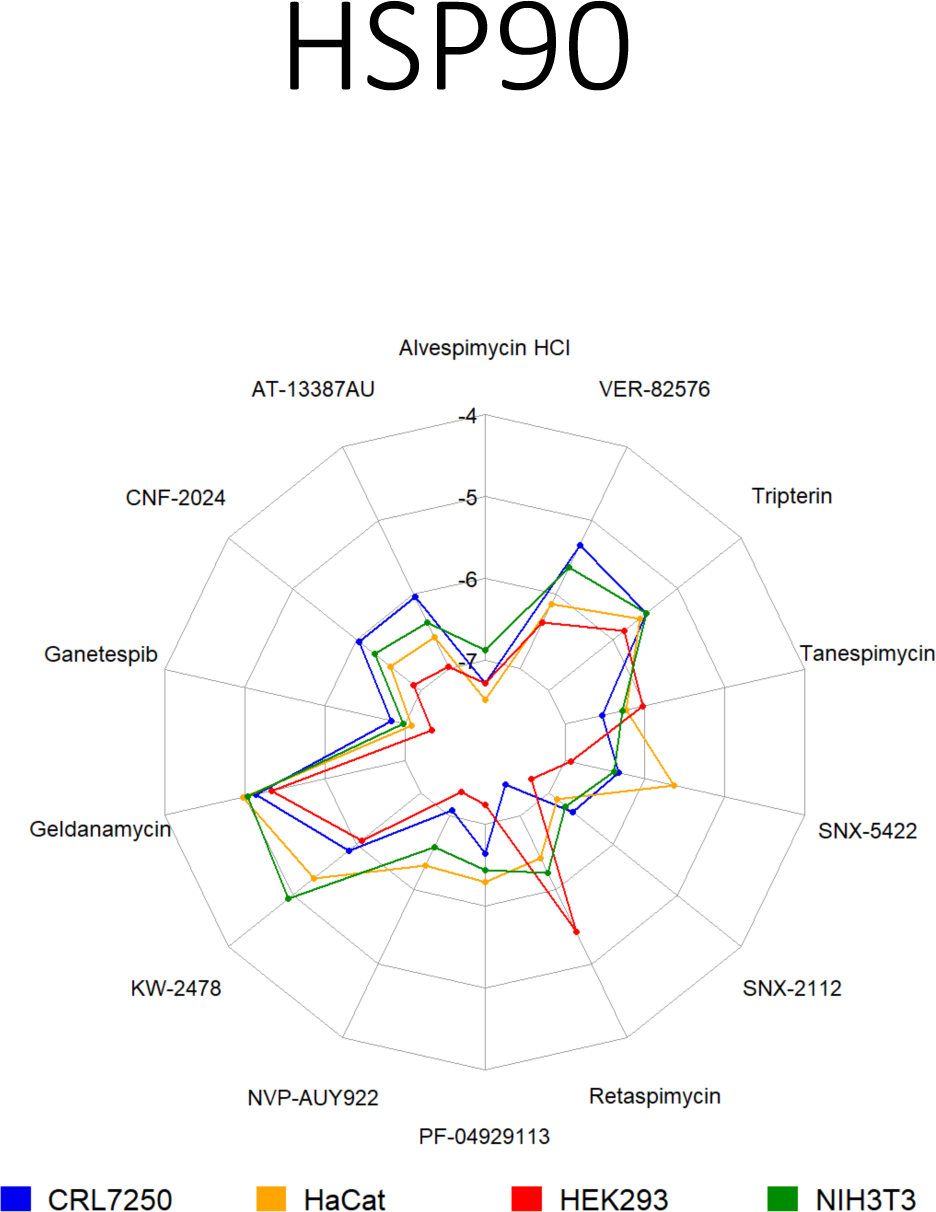
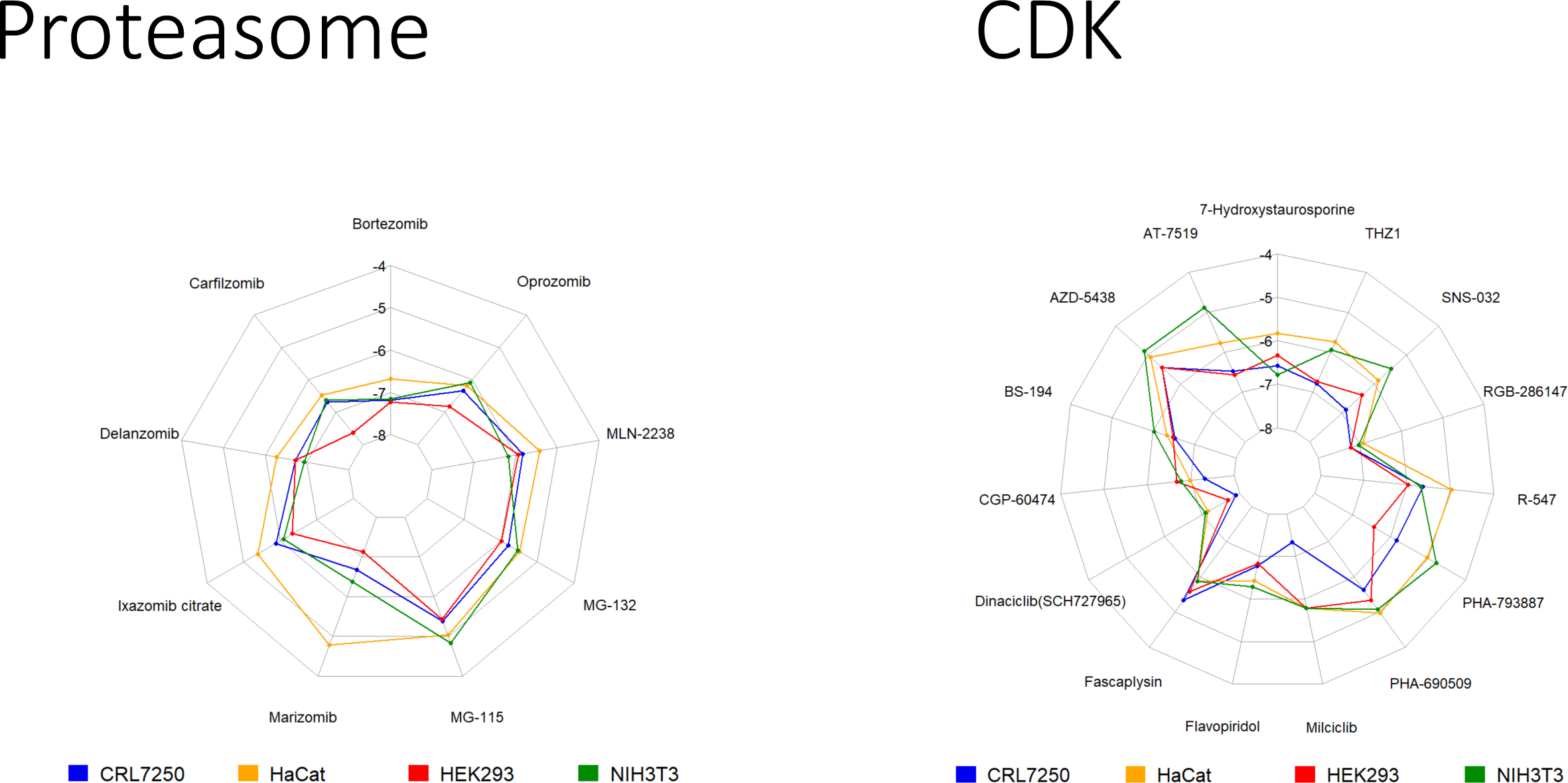
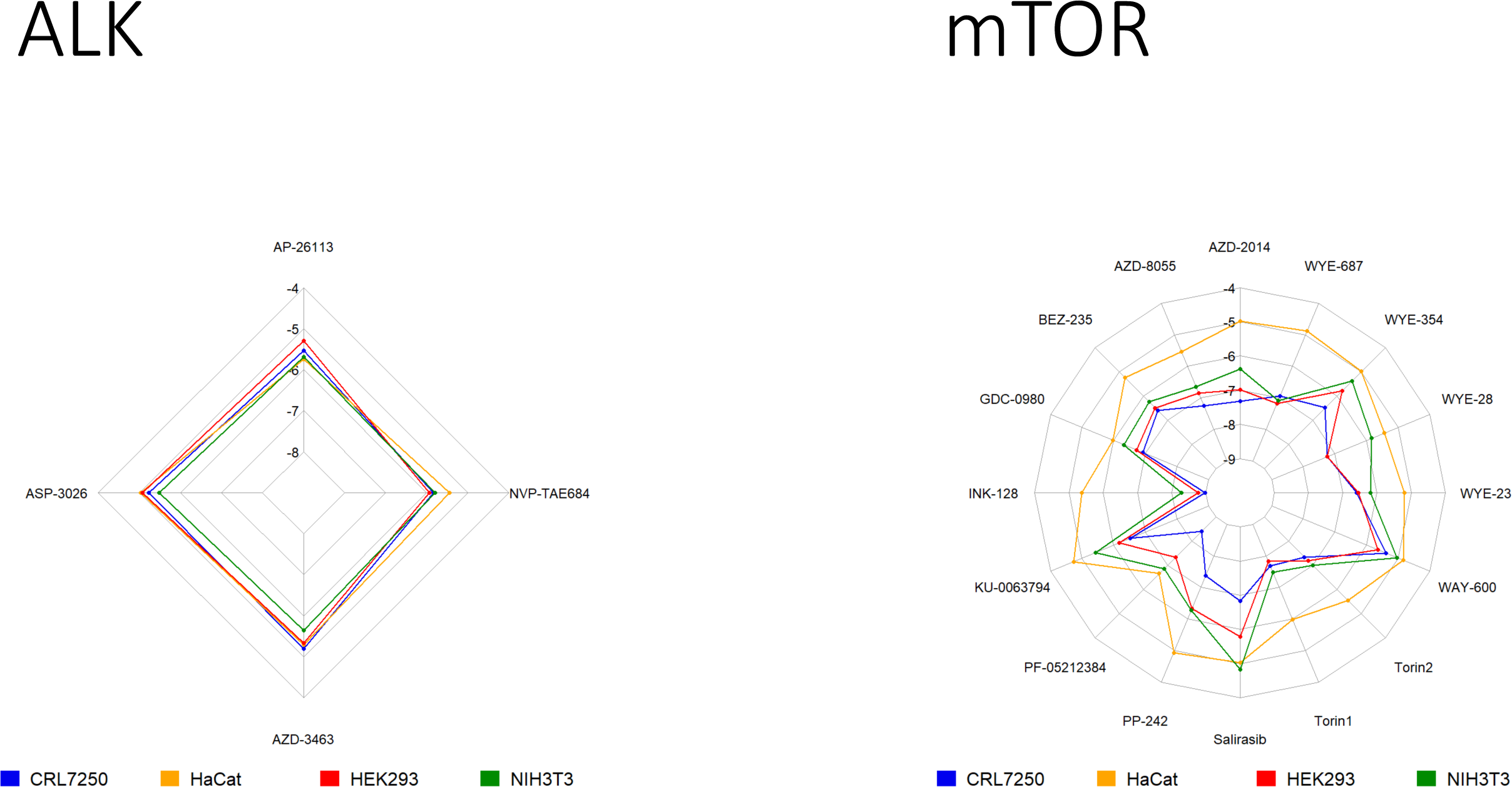
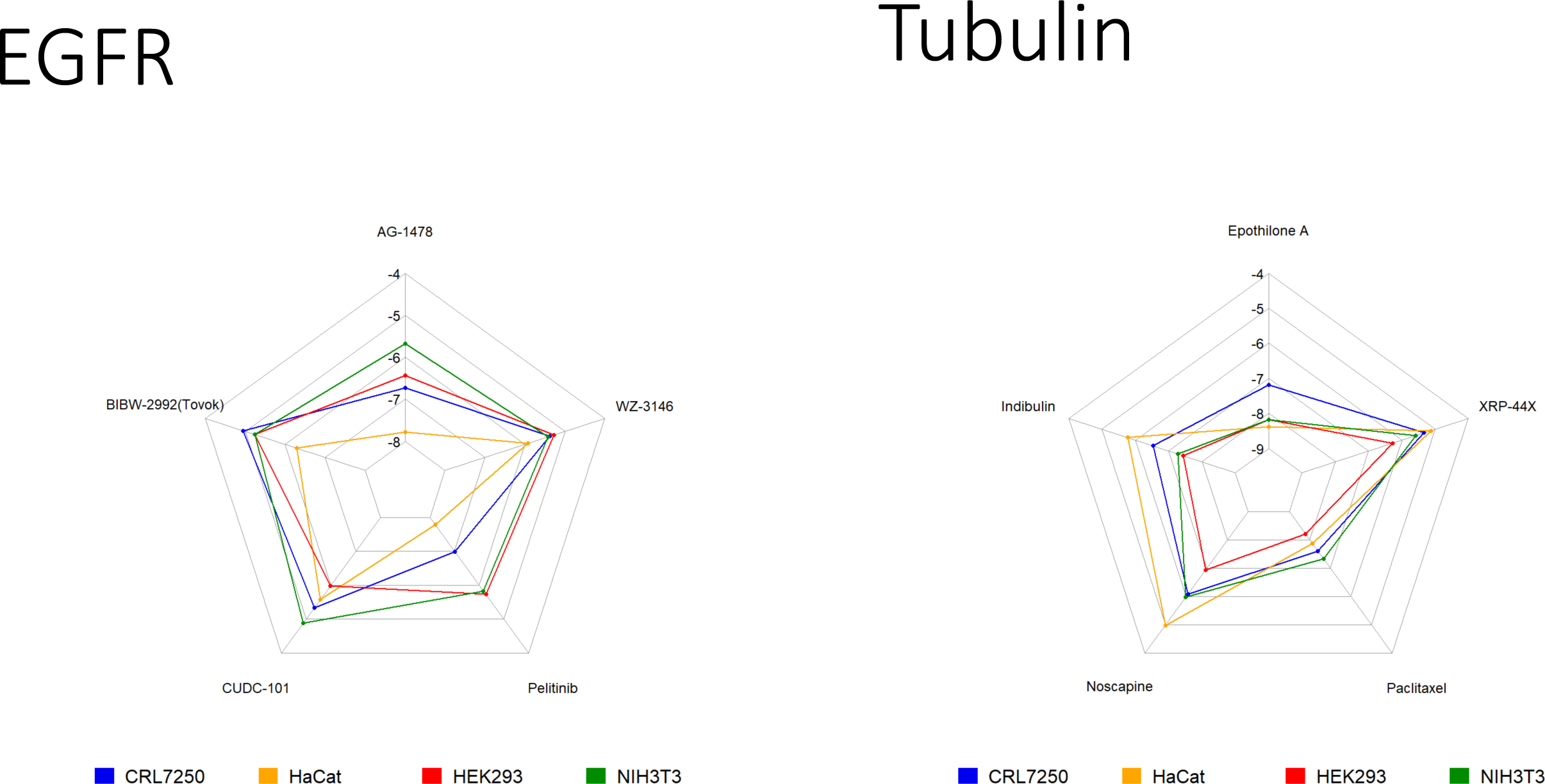
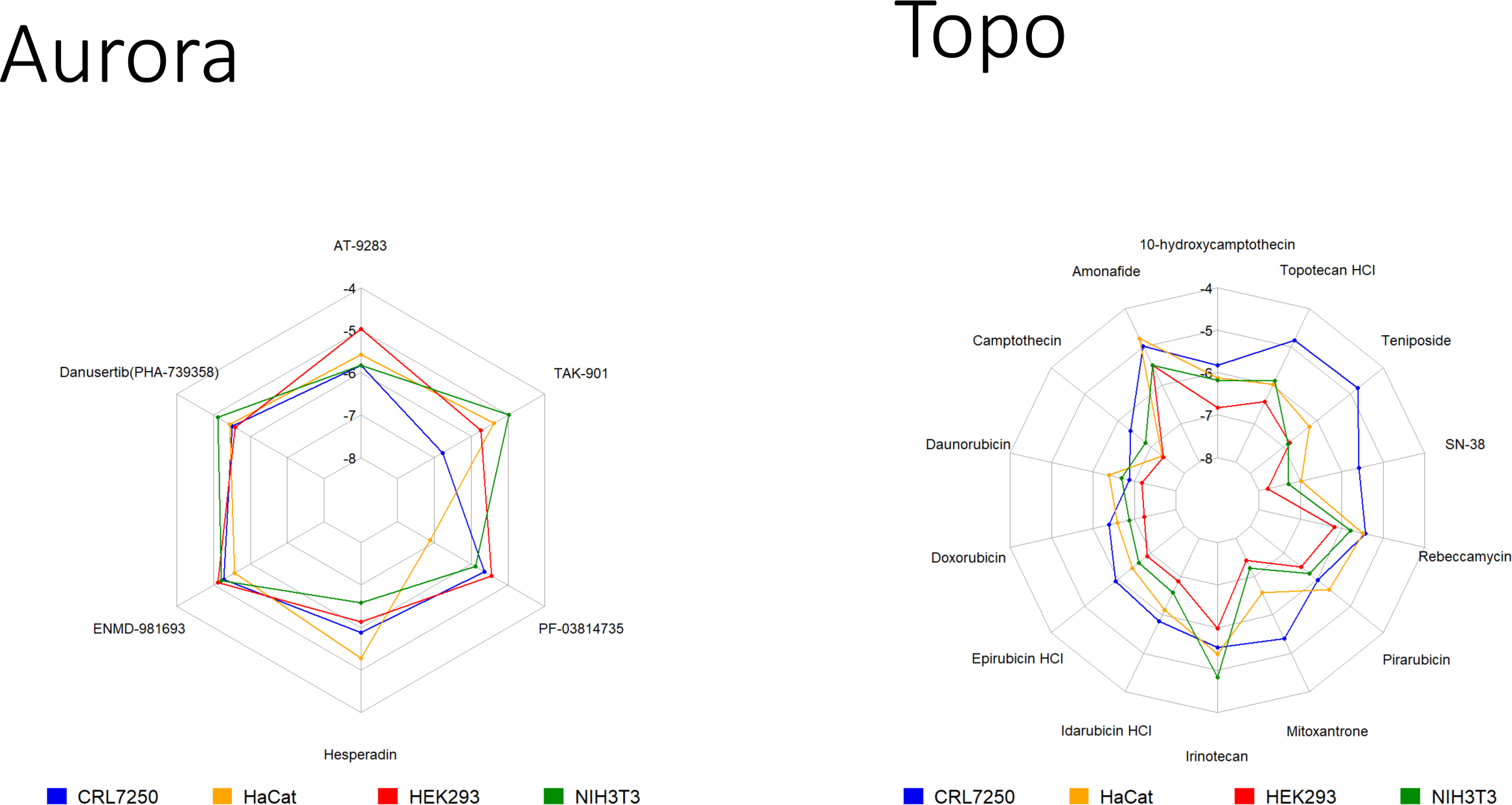
Radar plots displaying the Log EC_50_ for each compound in four normal cell lines. Radar plots are separated by MOAs that were overrepresented in the high-quality actives compounds such as HDAC, PI3K, HSP90, Proteasome, CDK, ALK, mTOR, EGFR, Tubulin, Aurora, and Topoisomerase.

**Supplementary Figure 2.**
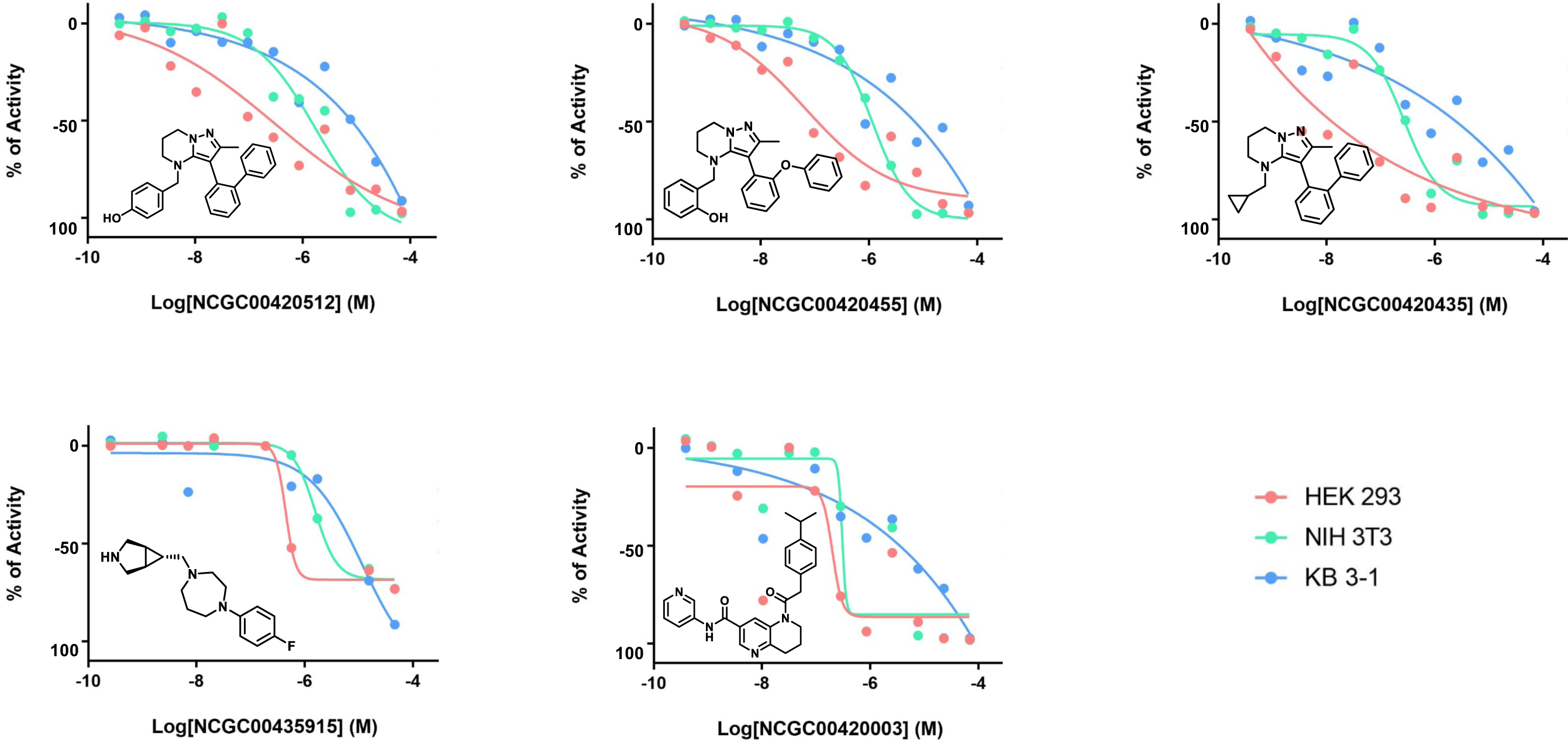
Dose-response curves for representative compounds in diversity library screen showing differential cell killing effect across three cell lines.

